# Commensal or pathogen: computationally vectorizing microbial genomes for *de novo* risk assessment and virulence feature discovery in *Klebsiella pneumoniae*

**DOI:** 10.1101/2024.05.13.593956

**Authors:** Kristen L. Beck, Akshay Agarwal, Alison Laufer Halpin, L. Clifford McDonald, Susannah L. McKay, Alyssa G. Kent, James H. Kaufman, Vandana Mukherjee, Christopher A. Elkins, Edward Seabolt

**Affiliations:** AI and Cognitive Software, IBM Research, 650 Harry Road, San Jose, 95120, CA, USA.; Division of Healthcare Quality Promotion, Centers for Disease Control, 1600 Clifton Road, Atlanta, 30329, GA, USA.; US Public Health Service, 1101 Wootton Parkway, Rockville, 20852, MD, USA.

**Keywords:** Klebsiella pneumoniae, virulence prediction, genomics, modeling

## Abstract

Bacterial pathogenicity has traditionally focused on gene-level content with experimentally-confirmed functional properties. Hence, significant inferences are made based on similarity to known pathotypes and DNA-based genomic subtyping for risk. Herein, we achieved *de novo* prediction of human virulence in *Klebsiella pneumoniae* by expanding known virulence genes with spatially proximal gene discoveries linked by functional domain architectures across all prokaryotes. This approach identified gene ontology functions not typically associated with virulence *sensu stricto*. By leveraging machine learning models with these expanded discoveries, public genomes were assessed for virulence prediction using categorizations derived from isolation sources captured in available metadata. Performance for *de novo* strain-level virulence prediction achieved 0.81 F1-Score. Virulence predictions using expanded “discovered” functional genetic content were superior to that restricted to extant virulence database content. Additionally, this approach highlighted the incongruence in relying on traditional phylogenetic subtyping for categorical inferences. Our approach represents an improved deconstruction of genome-scale datasets for functional predictions and risk assessment intended to advance public health surveillance of emerging pathogens.

## 1 Introduction

Virulence is the ability of a microbe to cause disease by inflicting damage upon the host [1, 2]. While this ability is observable and subject to Koch’s criteria, it becomes a difficult proposition to discreetly define at a molecular genetic level given a highly dynamic and contextual host-microbe relationship [3]. This has been exacerbated by the increasing wealth of genome sequencing that can blur the lines of typical gene content and plasticity within and between microbes of a given genus or species. They can differ genomically on a megabase scale due to acquisition of pathogenicity islands and other mobile content [4–6]. Also, noting that foods undoubtedly (and naturally) contain microbes obtained in food production chains, regulatory action is determined by virulence gene content that includes specific toxin subtypes and associated genetic accoutrement [7]. However, this targeting of specific virulence genes may miss emerging novel virulence factors or combinations thereof. Conversely, with beneficial microbes intended for consumption (e.g., probiotics), establishing a regulatory framework for assessing the safety of new strains to the market (apart from historical use) is not well-defined but rather relies on simple gene-level scrutiny based in part on available virulence database profiling (see Table 2 in Roe et al. [8]). Such capacity and risk assessment/scrutiny beyond antimicrobial resistance containment strategies is notably absent with healthcare-associated bacterial pathogens [9] but will undoubtedly require new computational approaches to move beyond simple gene attributes.

Virulence *sensu stricto* is generally defined as toxins and other directacting agents that form the basis of virulence and by extension includes factors that contribute to the adhesion, colonization, invasion, and immune-related evasion as secondary but important genetic repertoire [2]. However, manifesting virulence is a delicate interplay of the host-microbial relationship and ecology and is pleiotropic with effectors that include local and global regulatory elements, unique metabolic capacities, and utilization that may provide a competitive advantage within a given niche but are otherwise difficult to distinguish as part of the native (and required) genomic structural content [10]. A relevant example of this capacity is hypervirulent *Klebsiella pneumoniae* causing pyogenic liver abscesses and osteomyelitis in immunocompetent healthy individuals [11]. In this case, previously identified virulence markers were found to be associated with acquired plasmid content that included concerning carbapenemase-production and additional toxin-antitoxin systems of unknown consequence to disease manifestation albeit on separate virulence and resistance plasmids within the strain reported initially in the United States. Additionally, different independent multilocus sequence types (ST23 and ST11) provided the genomic backdrop permissive to such virulence and resistance convergence that has also further converged into single hybrid plasmid constructs [12–15]. Alternatively and from a broad context, integrative conjugative elements provide significant genomic plasticity from various cargo gene elements as an important nexus driving bacterial evolution with demonstrable ramifications for antimicrobial resistance gene representation [16].

The aforementioned analysis was aided by whole genomic sequencing datasets that provided an opportunity to connect with extant genes of interest/concern while highlighting additional factors that were associated with these elements. While the datasets provide a replete accounting of genetic content, we cannot advance our understanding, discovery, or synthesis of the interplay of related elements using a traditional gene-by-gene accounting. Genomics is poised to move these efforts into a prospective art similar, conceptually, to forecasting strain trajectories with predictive models for viral and other acute infectious diseases [17]. Some computational tools have leveraged machine learning with support vector machines (SVM) to identify and assess virulence such as VirulentPred [18] and VICMPred [19]; however, these implementations are often limited in scale and genomic diversity of the datasets thus limiting generalizability and accuracy in virulence forecasting.

Virulence databases, such as the Virulence Factors of Pathogenic Bacteria Database (VFDB) [20], exist for reference benchmarking of gene-level analysis and traditional scoring. However, considering that the associated factors are based on experimental and laboratory isolate analysis, they do not represent the full biodiversity of all microorganisms. Our central aim involved targeting functional deconstruction of relevant virulence factors into domain architectures through IBM’s Functional Genomics Platform (FGP) [21] that could be leveraged to discover hitherto unknown virulence-related proteins within publicly available genomes stratified by virulence categories. Once modeled with machine learning for inferring phenotypic virulence potential, model performance was evaluated for value-additions of the newly discovered virulence-associated proteins. We sought to address the virulence potential of *de novo* genome sequences of *Klebsiella pneumoniae*, a primary pathogen of significant consequence in healthcare settings and the environment [6]. Developing methods to expand our knowledge of virulence factors will improve our fundamental microbiological understanding and allow us to scale candidate biomarkers as organisms evolve, providing more robust data for public health tracking and surveillance.

## 2 Results

### 2.1 Prokaryotic virulence prediction method and scale

We developed a method that leverages gene co-localization and co-occurrence to identify putative virulence proteins augmenting existing references such as the Virulence Factor Database (VFDB) [20] by three orders of magnitude. This method was applied across 206,575 *de novo* and reference bacterial genomes spanning 1,400 genera in the Functional Genomics Platform (FGP) [21]. Key steps of our virulence prediction pipeline (Figure 1) relating to the scale of data analyzed are described here (full details in Methods Section 4.1 - 4.6).

**Fig. 1.**
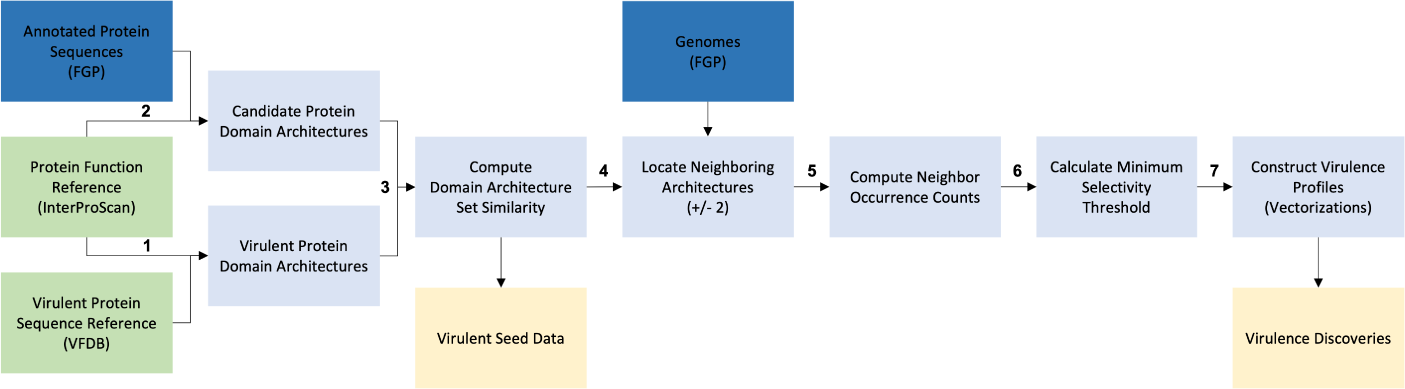
Virulent Phenotype Prediction Pipeline Schematic where data analysis steps (light blue), reference data (green), FGP data (dark blue), and key outputs (yellow) are shown.

Our method leveraged protein domain architectures (DAs) as its central feature representation, yet it is amenable to any coding system e.g., full gene or protein sequences. Domain architectures have been used for protein comparison [22–25] and for phylogeny [26] among others. In this work, InterPro (IPR) codes for 53M distinct protein sequences across 1,400 bacterial genera in FGP were collected to define the domain architectures. The resulting 297,004 distinct DA sets contained an average of 7.6 domains (median 5 domains) per DA (for distribution of domain types see Supplementary Table S1). Collectively, the DAs that represented all proteins in FGP were the candidate search space for our virulence prediction method.

Of the 28,583 known virulence factor proteins in VFDB [20] spanning 32 genera, 74% were an exact sequence match to an FGP protein and were annotated with a DA. To expand beyond exact protein sequence match, the set of DAs describing VFDB protein sequences were used to identify matching DAs within FGP proteins yielding 2,599 matching DAs which are collectively the “virulent seed data” (Table 1). All genomes analyzed contained at least one DA matching the virulent seed data. By leveraging domain architecture similarity, our method increased the number of virulent seed pivot proteins from 21,125 proteins in VFDB to over 11 million distinct protein sequences contained in FGP (Table 1) as well as expanded the number of included genera from 32 in VFDB to 1,400 in this analysis.

**Table 1.** Pivot Expansion and Neighbor Summary.

|  | Pivots | Neighbor Search Space | Discoveries |
| --- | --- | --- | --- |
| # DAs | 2,599 | 205,665 | 80,060 |
| # Protein | 11,144,804 | 32,723,557 | 16,278,692 |

Count values in this table represent number of DAs or number of distinct protein sequences for virulent seed data (pivots, P); co-pivots and potential discoveries prior to Kneedle analysis (neighbor search space); and candidate discovered virulence factors surviving Kneedle low frequency filter (discoveries, D).

During the FGP gene and protein annotation process [21], each gene/protein is identified with a relative position indicator serving as a location index within a genome. Genes are often co-located or clustered together when they participate biologically in support of a given function, and we exploited this to better construct virulence profiles.

DAs for virulent seed data are treated as “pivots” (P) to identify neighbors ±2 proteins— labeled as C for co-pivot (defined as a neighbor protein with a DA also present in the virulent seed data) or PD for a potential discovery candidate. This yielded a neighbor search space (C and PD) of 205,665 DAs and nearly 33M distinct protein sequences (Table 1).

Potential discovery candidates were removed using a low frequency filter with the Kneedle algorithm on a per-genus basis to result in the true discoveries (labeled as D). From this neighbor analysis, we identified 80,060 discovery (D) domain architectures from over 16M distinct protein sequences in association with known virulence factors.

Across the corpus of genomes analyzed, the number of unique pivot and co-pivot DAs per genome remained stable (mean = 750 and 1069, median = 780 and 998, relatively) with the number of discovery DAs observed at a slightly higher count (mean = 1528, median = 1555) (Supplementary Figure S1).

### 2.2 Discovered proteins associated with virulence across bacteria

In the standout discovery DAs (top twenty most occurring across all genera), ATP transport and transcriptional regulation were predominant in their enriched protein names/functions (Figure 2 and Supplementary Figure S2). These two molecular functions were also enriched when mapping to GO terms: GO:0055085 transmembrane transport and GO:0005524 ATP binding (Figure 2 and Supplementary Figure S2).

**Fig. 2.**
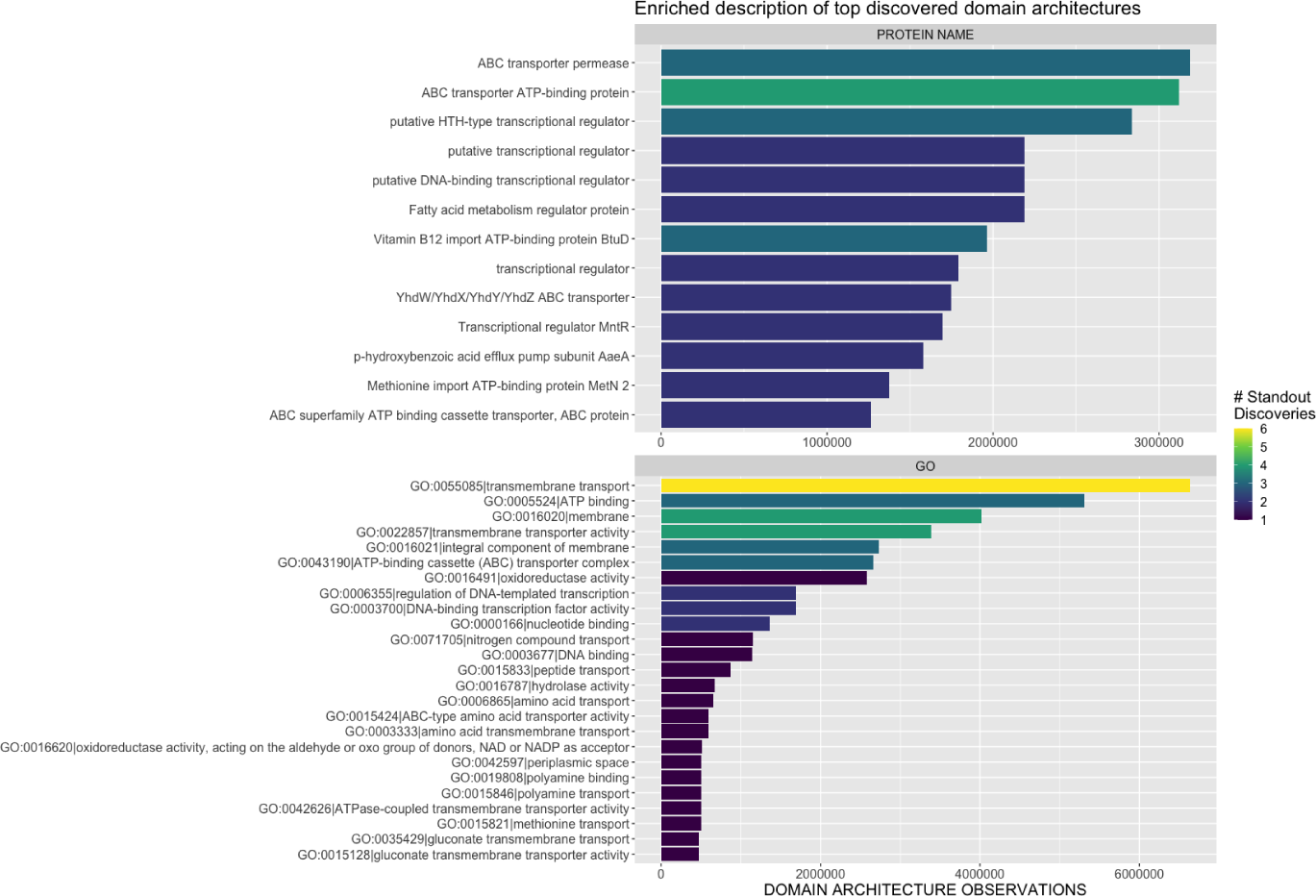
Enriched protein names and function for standout discoveries. The top 20 discovered domain architectures by highest occurrence were selected, and their respective enriched protein names (top panel) and GO terms (bottom panel) were identified. The number of top discovered DAs for each protein or GO term are indicated. For enriched protein names, the protein of origin was retrieved for each standout D DA. Protein synonyms (i.e. one domain architecture could be assigned multiple protein names) were common with 35 protein names on average per DA (range 6–83 protein names per DA). Therefore, enriched protein names (occurring for more than one top discovered DA and excluding “hypothetical” or “putative”) are plotted here. For the complete per DA breakdown, see Supplementary Figure S2.

Standout discoveries were observed to appear with a diverse multitude of pivot proteins; on average, each standout discovery was observed with 411 different pivot DAs (range 130–694) indicating promiscuity of pairing.

Relating to pairing frequencies, we calculated the relative occurrence of pivots within the genome cohort (as a percentage) and the percentage of the neighbor to yield 799,552 pivot:discovery pairings and 270,525 pivot:co-pivot pairings (Supplementary Figure S3). Pivot DAs associated with discoveries occur in higher copies across the genome cohort than with co-pivot neighbors (Supplementary Figure S3). This is commensurate with previous findings of increased copy number indicating increased virulence in *Yersinia* [27], *Salmonella* [28], and *Renibacterium* [29].

Using k-means clustering, we selected pairings with the highest relative occurrence (cluster 4 for pivot:discovery and cluster 6 for pivot:co-pivot pairings) and mapped neighbor DAs to their respective GO terms. There was 94.4% overlap in GO terms between the pivot:co-pivot and pivot:discovery enriched clusters indicating significant functional similarity with top terms such as serine-type endopeptidase activity (GO:0004252), ATP binding (GO:0005524), DNA binding (GO:0003677), metalloendopeptidase activity (GO:0004222), and regulation of DNA-templated transcription (GO:0006355). Beyond this, there were 2,529 GO terms unique to the pivot:discovery enriched cluster with RNA binding (GO:0003723), aminoacyl-tRNA ligase activity (GO:0004812), translation (GO:0006412), and tRNA aminoacylation for protein translation (GO:0006418) being the most frequent terms. There were only 21 GO terms unique to the pivot:co-pivot enriched cluster (all with low frequency *<* 5). The high level of functional similarity in these enriched clusters further supported the inclusion of discoveries as putative virulence factors, while the additional functional terms aid in providing a comprehensive description of virulence molecular function and biological processes.

### 2.3 Case study: machine learning prediction of *Klebsiella pneumoniae* virulence

To further assess the utility of these virulence discoveries, we completed a case study on *Klebsiella* with an application of machine learning to stratify and predict virulence risk.

We defined four virulence classes based on subject matter expertise leveraging the sample’s isolation source (Supplementary Table S2). In summary, samples collected from more sterile sites e.g., blood, aspirate, cerebral spinal fluid are considered highly virulent. Samples collected from non-sterile body sites e.g., drainage, sputum, respiratory are considered low virulent. Colonization samples are defined as samples originating from rectal swabs, skin, or feces. Samples labeled as not virulent samples were isolated from environmental or animal samples.

The domain architectures for 2,802 *Klebsiella* genomes were used for two model comparisons: 1. high versus low virulent and 2. virulent versus not virulent (Methods Section 4.8) representing risk prediction needs for public health. Our best performing model for classifying high versus low virulent samples achieved an F1-Score of 0.81 with an AUC of 0.82 with a Class1 precision of 0.84 for the high virulence phenotype (confusion matrix in Supplementary Figure S4a). For virulent versus not virulent samples, our best performing model achieved an F1-Score of 0.80 (confusion matrix in Supplementary Figure S4b). To contrast the usage of our virulence discoveries versus current published data, we evaluated baseline models with only domain architectures derived from VFDB virulent seed data (Figure 3a). For these best performing models, inclusion of our virulence discoveries yielded a 2% and 4% improvement, respectively. Across all sixteen paired model evaluations, discovery inclusion improved the F1-Score or achieved parity (within 0.01) with 7.5% increase being the largest improvement. Additionally, the Class1 precision and Class0 recall were higher when including our putative virulence discovery features (Figure 3b). It is known that when the number of features far surpasses the number of samples that machine learning model accuracy can be dampened [30]. We suspect this to affect the discovery inclusion models which originated from 3,905 features (compared to the 1,193 features for baseline). As genomic surveillance continues to expand, the number of available public genomes will increase as well and will undoubtedly allow for model improvement and refinement.

**Fig. 3.**
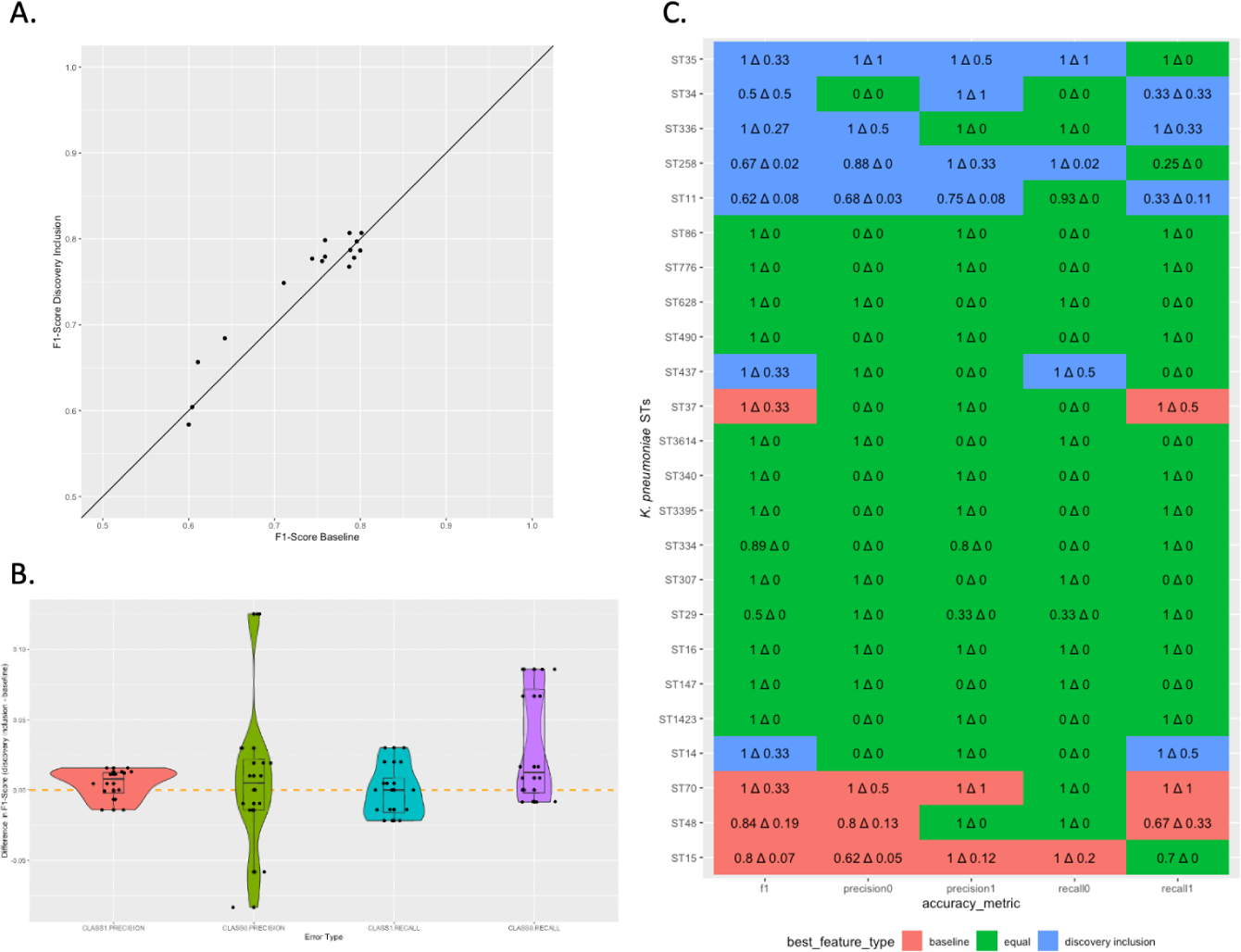
Model performance on virulence prediction. Virulence prediction models are compared when using baseline data based on VFDB virulence seed data alone (baseline) and for our discovery inclusion features with regard to the model’s F1-Score (**A**) and for the class precision and recall (**B**). Two comparisons were modeled: virulent versus not virulent and high virulent versus low virulent. In each model comparison, Class1 denotes the more virulent sub-group and Class0 denotes the less virulent sub-group. In **B**, the difference of our discovery inclusion minus the baseline score is shown where positive values indicate an improvement from the discovery inclusion feature set over baseline features. Error analysis was completed and is shown for MLSTs with greater than one genome in the test set. (**C**) shows the best performing feature set per ST with the value per accuracy metric shown with the delta (difference) of discovery inclusion and baseline.

Model error analysis was completed per multilocus sequence type (MLST) to assess the limit of classification accuracy by lineage. Across all models, the baseline and discovery inclusion feature sets performed comparably with respect to F1-Score, precision, and recall by class (Supplementary Figure S5). Discovery inclusion models performed slightly better in the case of MLSTs with fewer genomes in the test set (Supplementary Figure S5a). To represent real-world usage of these models, we further examined models with the highest F1-Score (independent of error analysis outcomes) for each classification task: virulent versus not virulent and high virulent versus low virulent. When classifying virulent versus not virulent samples, the discovery inclusion and baseline feature sets performed comparably; however, when comparing high versus low virulence, the discovery inclusion feature set yielded higher F1-Scores, precision, and recall for each class (Supplementary Figure 5c). This indicates that expanding the feature set provides additional ability to discriminate between samples that may be more similar in their virulence risk. Specifically, the virulence potential in *Klebsiella pneumoniae* (Kp) ST35, ST34, ST336, ST258, ST11, ST437, and ST14 were all better predicted using the discovery inclusion feature set. The virulence category for Kp ST37, ST70, ST40, and ST15 were better predicted using the baseline feature set.

Since assessment of virulence and phenotypic risk is often still assessed based on taxonomy, we completed MLST analysis of the isolates and contrasted those against virulence category. Sequence type alone did not separate the virulence classes (Figure 4). For example, of the 38 most common STs, 35% were assigned to more than one virulence category, and 74% were assigned to more than two virulence categories. We did observe an increase in diversity of MLST in the colonization and not virulent categories compared to high or low virulence categories (Figure 4 and Supplementary Table S3).

**Fig. 4.**
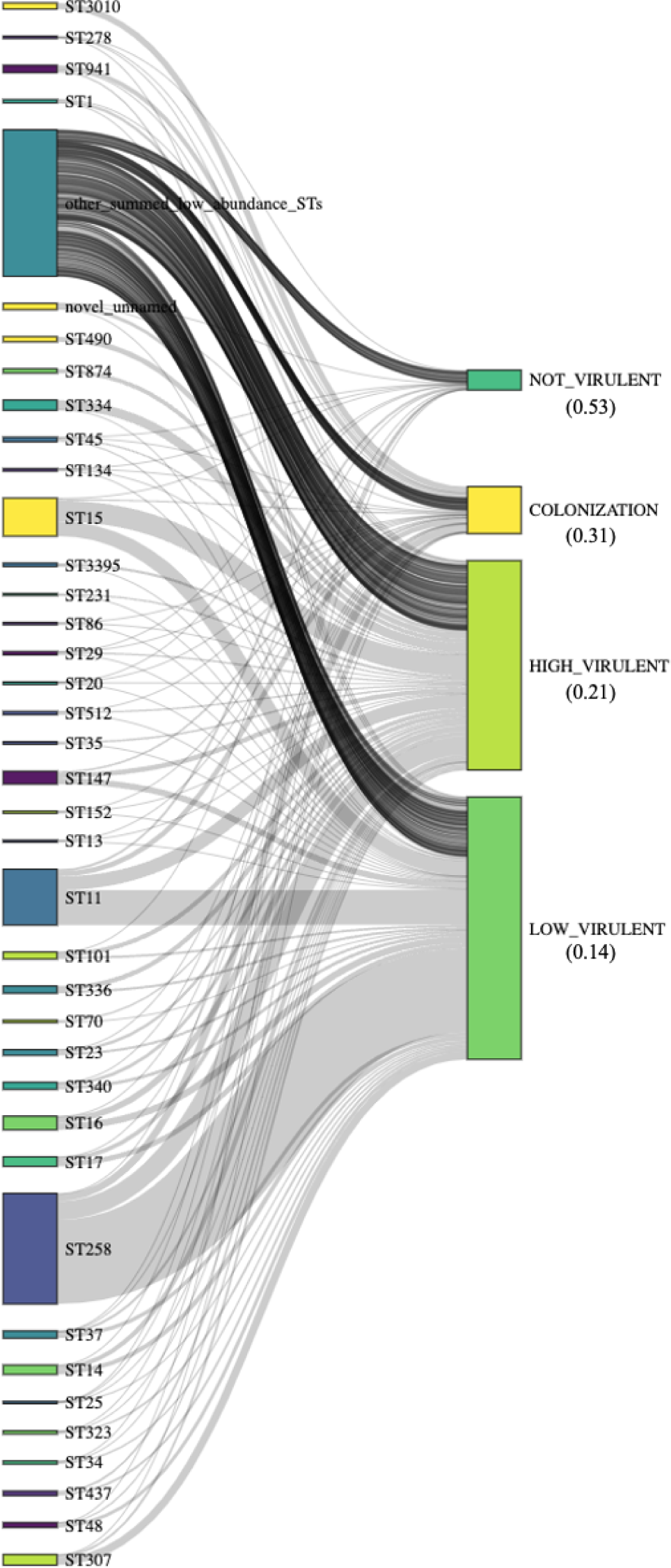
Comparison of top *K. pneumoniae* isolate MLSTs with virulence category. STs are indicated using the Pasteur scheme *Klebsiella pneumoniae*. MLSTs with less than ten genomes are binned as other low abundance STs. Virulence categories are ordered by relative MLST diversity per category (shown in parentheses beneath the category label) which is defined as number of MLST per number of samples for each virulence category (full counts are shown in Supplementary Table S3.

In addition, we leveraged a marker gene approach, Kleborate [31], to complete a virulence score analysis and contrasted these scores against our virulence categories (Figure 5). Kleborate focuses on the virulence loci yersiniabactin, colibactin, and aerobactin to define a virulence score 0 – 5, with increasing score indicating an increase in virulence severity. We observed a mixture of virulence scores from Kleborate across our labeled virulence categories where increasing virulence severity did not align with an increase in score (Figure 5a). For the highest Kleborate virulence score, we observed a comparable proportion of isolates 0.5% – 1.2% per virulence category of colonization, high or low virulent. Low virulent isolates were observed to have a higher proportion of score 1, presence of yersiniabactin only, and more score 2, presence of yersiniabactin and colibactin or colibactin only than other virulent categories. Colonization and high virulent categories were observed to have a comparable proportion of score 4 aerobactin with yersiniabactin (no colibactin) isolates, 3.2% and 3.1% for colonization and high virulent isolates, respectively. Isolates labeled as not virulent did exhibit a higher percentage of virulent score 0 with no virulent loci detected.

**Fig. 5.**
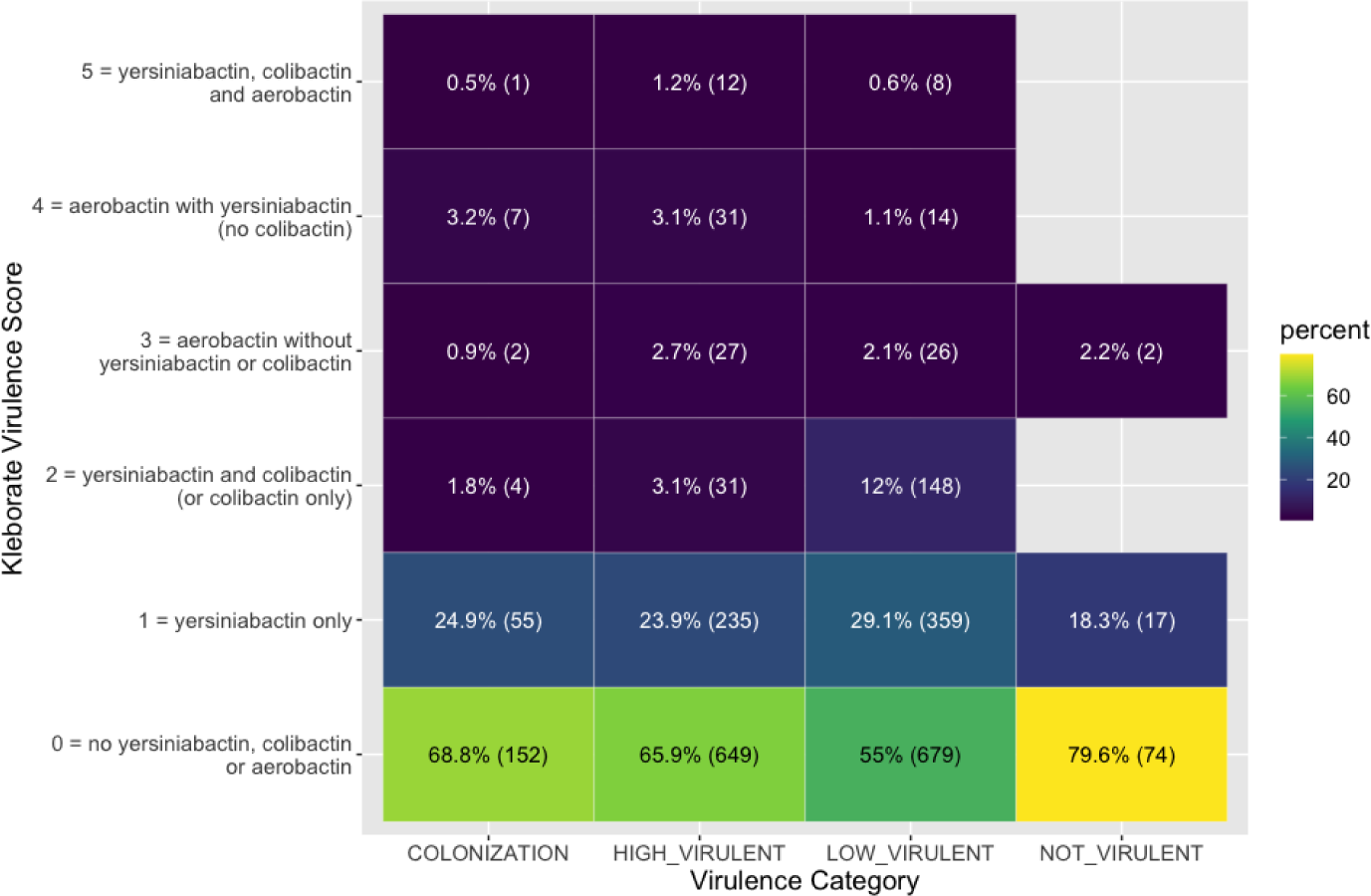
The virulence category for *Klebsiella* isolates is contrasted with the virulence score as identified from Kleborate. The percent of isolates per virulence category is shown (i.e. 55% of low virulent isolates exhibited a Kleborate virulence score of 0) as well as the count of isolates per category set.)

## 3 Discussion

We successfully developed a rational approach to utilize expansive public genomic datasets to better predict virulence potential in *Klebsiella pneumoniae*. A strength of this approach is that it was based upon genera-agnostic discovery of additional putative virulence factors using the entire VFDB for seed data with performance that extends beyond clinical severity phenotype-specific methods such as Kleborate [31]. Therefore, the application and model development can be moved into any taxonomic strata using the framework we established. Incorporating such approaches to iteratively refine model development while acknowledging that log order increases in magnitude of genomic data will undoubtedly ensue, and enhance/increase domain architecture seeds and, relatedly, markers of virulence.

Koch’s molecular postulates provide a rationally defined, albeit restrictive, basis for identifying gene-specific contributions to disease by a pathogen [32, 33]. Accordingly, traditional laboratory-based approaches to virulence assessment have been an important pursuit to discriminate particularly troublesome strains, characterize underlying molecular mechanisms, and identify specific, robust biomarkers for detection and screening assays [34]. While successful, these are typically focused on specific severe clinical disease phenotypes and experimentally-defined structural genes with crucial, rationally-defined involvement in virulence processes. Relatedly, virulence databases, such as the VFDB employed herein, place a significant emphasis on such characteristics for database inclusion. Through the course of our work, we were able to expand on the restrictive boundaries of such approaches, with less intuitive and more nuanced genetic content/associations attributable to spatial relationships and their functional roles through protein domain architectures. The molecular architecture of bacterial genomes has spatial relationships due to selection pressure, as spatially organized genes related to a common function (i.e., operon) allows for co-regulation and improved/efficient response to rapid changes in the environment [35]. Furthermore, segregation of encoded “accessory genome content”, such as virulence factors, coalesced over time on islands or prophage remnants, as has been well-documented [35–37], allowing for these traits to traffic together or be selected out.

In the discovery phase of our approach, several new putative virulence-associated genes were uncovered. Expectedly so, these were enriched for membrane-associated attributes such as transport and ATP-binding cassette proteins. In addition, gene regulation-associated functions which are less intuitive from a functional perspective without specific experimental validation, are usually identified with time- and resource-intensive studies in animal models of virulence [38]. Hence, gene ontologies and associated functions with discoveries herein can have many pleiotropic effects either with their experimentally derived molecular genetic manipulation through overexpression, deletion, and complementation that yield less than clearly defined functional consequences. Tangentially, an example of these difficult-to-identify relationships with virulence is exemplified with tellurite resistance in many O157:H7 and non-O157 serotypes of Shiga-toxin-producing *E. coli* (STEC). While this characteristic has a notable association with the STEC pathotypes, its relationship to virulence *per se* and distribution in other genera is still unclear, but it has spatial relationships with a variety of other bacterial functions [39]. Relatedly, in our study with *Klebsiella*, the findings are commensurate with a body of literature associating tellurite resistance with pathogens with a classical basis for clinical microbiology selective culturing [40, 41]. Hence, several domain architectures map to proteins names associated with “tellurite” and included 3 pivots and 13 discoveries across all genera, although tellurite GO terms were not mapped with our domain architecture-associated protein names. Nonetheless, these 3 conserved tellurite-associated domain architecture pivots and 2 of 13 discovered domain architectures were found widely dispersed in nearly all *Klebsiella* strains (5,680 and 5,658, respectively, of 5,689 genomes). For future analysis, it expands the feature set to identify tellurite resistance and, by extension, pathogens that may be resistant. It is also noteworthy that discoveries occur with a high diversity of pivots, indicating there is promiscuity of pairing between the discoveries and pivot proteins. This observation would argue that nascent discoveries have a supportive connection with virulence in general and provides for their legitimacy as opposed to being simply a surrogate for a specific virulence factor (pivot).

For our purposes, we were able to test the strength of these discoveries by measuring the performance improvement of the resultant models, wherein they were folded into the analysis for measurable value-added context. Compared with baseline measurement using only VFDB seed data, the models achieved as much as 7.5% improvement in F1-Score in test datasets. In an analogous framework, compared to another available virulence scoring system, Kleborate [31], virulence severity exhibited mixed associations with our classifications. However, given the limited stratifications that can be achieved with Kleborate based on only a few biomarkers of virulence versus the pangenome scale of our approach, the comparison is highly uneven and difficult to interpret beyond these observational findings which is noteworthy to perform given the limited tools currently available.

These classification models were also assessed for traditional MLST schemes. Considering that the study scoped all publicly available *Klebsiella* genomes, we did not consider the MLST assignment when the virulence category definitions were created. So, it is interesting then to observe resultant MLST diversity by virulence category with “Not-Virulent” and “Colonization” encompassing higher indices than “High- and Low-Virulent” classifications (Supplementary Table S3). For example, in the 38 most common MLSTs, 35% were assigned to more than one virulence category, and 74% were assigned to more than two virulence categories. Following this logic, our findings support that there are functional features beyond phylogeny influencing the virulence potential of an organism. This has practical importance and interpretation given reports of genera-specific commensal diversity as opposed to the emergence (and evolutionary convergence) of epidemic clones of respective pathogenic genera. This has been documented notably in *E. coli* [34, 42] and recently highlighted a targeted need for *Klebsiella* [43]. It is noteworthy as well that backend analysis of unique features contributing to model performance in *Klebsiella* contained significant numbers of hypothetical proteins (Supplementary File S2) that may further advance functional discoveries.

While this work is computationally intensive on the front-end for model development and refinement, the resulting models demonstrably improved our ability to generate a completely *in silico* assessment of respective genomes in this regard. However, it is an important caveat that computationally-derived dispositions of respective genomes are inherently based on key metadata features, attributing source as a primary driver for such categorizations in the ability to identify, resolve, and properly recall virulence categorizations *de novo*. Interestingly, this limitation underscores an important need for public repository metadata standards [44] but also in further instructing and refining relevant metadata that can assist and expand on computational applications to advance our understanding of microbiology at log orders higher analytical magnitude. In addition to improved metadata, future curated data sets of WGS are needed that are more representative of the genetic diversity across various epidemiologic categories informative of predictive models for human virulence (i.e. environmental, animal, and human isolates, including colonization, low and high clinical virulence/invasiveness). Thereby, as an extension, this approach can theoretically scale to any feature database in an exploratory manner for interrogating specific functional potentials for genomes of interest even beyond virulence or alternatively tailored/tuned to specific virulence subclassifications/manifestations.

Finally, an important limitation of this study and pathogen-associated genomic interrogations in general, is that they do not account for host factors contributing to disease severity (including host genotype) and clinical presentation [1]. Examples include methicillin-resistant *Staphylococcus aureus*, hypervirulent *Klebsiella pneumonia*, and enterohemorrhagic *E. coli* capable of causing hemolytic uremic syndrome (HUS). Furthermore, *E. coli* O104:H4 had a propensity for HUS development more so in females [45, 46]. Noting in the ensuing genomic analysis of the outbreak strains that traditional typing does little to provide insight into fundamental information about a strain’s virulence attributes [47]. Relatedly, metadata used to capture virulence categorization/disposition is only a snapshot in time of potentially significant differences and pathways of disease progression or amelioration, and even environmental attribution. Having avenues to provide additional context or clinical endpoints would undoubtedly refine model development and associated performance. Ultimately, incorporating virulence in national bacterial surveillance is a critical progression to leverage the increasing genomic data wealth, especially with rational approaches described herein. Relying on traditional phylogenies and typing schemes [5] do not align well with disease potential and only in certain already emerged epidemic clones with well-described virulence attributes can action be driven based solely on clonal structures. Approaches such as these, may move surveillance from a retrospective temporal art to predictive forecasting to identify pathogens of concern early in their emergence.

## 4 Methods

Our bacterial virulence predication pipeline methodology is depicted in Figure 1. The compute environment consisted of a single bare metal machine having 56 Intel(R) Xeon(R) CPU E5-2690 v4 @ 2.60GHz, 250 GB RAM, and running Ubuntu 20.04. Python 3.9.10 and Apache Spark 3.2.1 were used for implementing our prediction pipeline and analysis of data. Python 3.10 was used for the machine learning use case.

### 4.1 Annotated protein sequences

Protein sequences and their corresponding protein domains were retrieved from the Functional Genomics Platform (FGP) annotated genomes. To accomplish this, these sequences were derived from bacterial isolates spanning 1,400 genera and comprised a total 206,575 genome isolates (*de novo* genome assemblies constructed as previously described in Seabolt et al [21] and assembled genomes from NCBI RefSeq and GenBank). Each genome fasta file contained the largest contiguous scaffold and shorter DNA sequences that could not be integrated into other contigs or scaffolds and which may represent plasmid sequences. Genome assemblies were annotated using the FGP pipeline to identify their genes, proteins, protein domains, and corresponding function as described in Seabolt et al [21].

For domain identification, we leveraged InterProScan which uses thirteen protein signature databases e.g., CATH, Pfam, CDD to ascribe InterPro (IPR) domain codes to provide the known function of proteins. Of 53,836,264 distinct protein sequences in FGP, 42,724,008 (79.4%) proteins were annotated with at least one domain (IPR code) and are shown with their IPR type (e.g. DOMAIN, FAMILY, HOMOLOGOUS SUPERFAMILY, BINDING SITE) in Supplementary Table S1. The data representing these sequences were imported into our Apache Spark instance as parqueted tables (a column-based data storage format for performance optimization).

### 4.2 Protein function reference

For each protein sequence in FGP, the set of unique IPR codes were ordered alphanumerically to define the Domain Architecture (DA). In total, creating a collection of 297,004 distinct sets. For each DA, we computed the MD5 hash of the IPR code set to simplify query operations across tables. Collectively, the DAs that represent these proteins are the candidate protein search space for our method. See Supplementary Table S1 for a summary of the IPR codes and proteins in FGP.

### 4.3 Virulent protein sequence reference and domain architecture similarity calculation

Known virulent protein sequences were obtained from the Virulence Factor Database (VFDB) [20]. The VFDB protein dataset (retrieved December 2019) was parsed and persisted in our Apache Spark instance as a parqueted table (see Supplementary Table S4 for summary). This consisted of 28,583 distinct protein sequences from VFDB across 32 genera where 74% were an exact sequence match to an FGP protein sequence and were annotated with a DA. Each protein sequence was hashed using MD5 to create a unique string representation of that sequence to facilitate query operations across the different tables of data used in our pipeline. To expand beyond merely an exact sequence match, the DAs from VFDB protein sequences were also used to identify protein sequences within FGP with matching DAs. Similar domain architectures were defined as an exact match of all member IPR codes between two proteins representing 2,599 DAs which are collectively the “virulent seed data” with known virulent phenotypes.

### 4.4 Locating neighboring domain architectures

During the gene and protein annotation process [21], each gene/protein is identified with a relative position indicator within a given genome where numerical proximity indicates genomic proximity. We use this relative location index to identify neighbors within a genome.

With our virulent seed data and the FGP genomes, we constructed a table that records the NCBI accession number, DA, and location index of the virulent DA within a genome. These DAs are called “pivots” relating to neighbor identification and their known virulence function (Methods Section 4.3). Once the pivots are known, neighbors are identified that match the genome accession number and location index within a distance of ±2 proteins from the pivot. Neighbors are labeled as either P for pivot, C for co-pivot (a neighbor protein with a DA present in the virulent seed data), or PD for a potential discovery candidate (i.e., a protein discovered to potentially be associated with virulence). Refinement of discovery candidates to true discoveries are described in Methods Section 4.5. P DAs will always have a 0 normalized location index. C DAs will always be a pivot protein found in the pivot set, but will co-occur as a neighbor to another P DA.

Protein neighbors without an annotated DA are removed in a filtering step for quality control as they could indicate the end of the contig or scaffold, or could be due to missing data.

### 4.5 Computing neighbor occurrence counts

Neighbor occurrence counts were calculated per genus from the neighbor analysis performed in Methods Section 4.4. These counts are used to construct a dynamic high pass filter for removing low frequency neighbors aiding in the construction of a final table best representing virulence profiles with the highest concentration of co-occurring DAs with associated virulence. Low frequency neighbors were removed by calculating a minimum cutoff threshold using Kneed [48], an open source Python package implementing the Kneedle algorithm which works by utilizing the per genus neighbor count data sorted in descending order and returning the knee point of the fitted function to this data. Here the minimum cutoff threshold is represented by the y-axis value of the knee point. Any neighbors that were marked as potential discoveries are now considered true discoveries if they survive the filter step. This package was implemented as an Apache Spark User Defined Function (UDF) and computed at scale for all bacterial genera in FGP. Supplementary Figure S6 shows an example of this automated dynamic selection of the minimum threshold for *Klebsiella*.

### 4.6 Virulence profile construction and vectorization

Using the finalized neighbor counts from Methods Section 4.5, vectors of the predicted virulence profiles were created to generate two representations. First, for each genus and P DA, vectors of the neighbors were created using the following format:

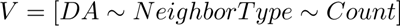

where *V* is the vector, *DA* is the MD5 hash of the neighbor DA, *Neighbor Type* is C for co-pivot or D for discovery, and *Count* is the number of times the neighbor was observed.

Secondly, for each genome accession number, vectors of the DAs were generated using the same format as the first. However in this representation, all DA types are included (P, C, or D) with the corresponding DA *Count* by type.

### 4.7 Site based domain bias assessment

IPR terms consist of multiple types including standard protein domains: DOMAIN, FAMILY, HOMOLOGOUS SUPERFAMILY which each will span a sub-chain of amino acids in a protein, but also types that are more localized to a single or very short amino acid region. These latter type of terms include ACTIVE SITE, BINDING SITE, CONSERVED SITE, PTM (posttranslational modification), and REPEAT. Since these denote as few as one amino acid and could potentially inflate count-based analyses, these site-based terms were assessed for their occurrence in relation to all domain architectures (Supplementary Figure S7) and within discovered domain architectures across all genera studied (Supplementary Figure S8). Of the 297,004 unique domain architectures in the seed data only 843 DAs were comprised exclusively of site-based terms, and across all domain architectures on average only 12.2% of member terms are site-based (median 5.4% site-based terms). The prevalence of exclusively site-based domain architectures in our D, P, and C domain architectures was found to be present in less than 0.2% of DAs with discovered DAs exhibiting less exclusively site-based DAs than pivot or co-pivot DAs (Supplementary Figure S8). In comparison, on average 99.6% of DAs per genus were intermixed with standard domain and site-based terms. Together, this indicated that we were unlikely to be affected by inflated counts based on low information content from site-based terms and therefore do not need to normalize or remove these domain types.

### 4.8 Virulence prediction using traditional and machine learning methods in clinical *Klebsiella*

#### 4.8.1 *Klebsiella* data description

To assess the discovered domain architectures for their utility in predicting virulence, we used these data hereto referred as “discovery inclusion” as a feature set for prediction of multiple virulence classes relating to clinical *Klebsiella*.

First, genomes labeled *Klebsiella* (NCBI Taxonomic ID: 570) were identified within FGP. This *Klebsiella* cohort consisted of 5,689 genomes of which 5,186 are of the species *Klebsiella pneumoniae* (Kp). MLST were assigned as described in Methods Section 4.8.2. The genomes that were not Kp consisted of 251 *K. aerogenes*, 238 *K. oxytoca*, 13 of unknown species, and 1 *Enterobacter cloacae*. Thus, approximately 10% of this cohort were non-Kp genomes. For this work, isolation source was used heavily in our virulence class definitions and therefore, was considered required metadata for each genome. If the isolation source was absent or ambiguous e.g., isolation source contained nondescript information as in “sterile body site” or “United States” instead of a well-defined isolation source e.g. “feces” or “blood”, then the genome with the incomplete isolation source information was removed from this cohort. This resulted in 2,802 *Klebsiella* genomes from multiple isolation sources including 2,518 from human hosts of varying body sites.

#### 4.8.2 Application of traditional bioinformatic methods in *Klebsiella* use case: k-means clustering, taxonomic, and virulence characterization

Unsupervised clustering of the discovery inclusion feature set was performed using k-means and DBSCAN; no clusters were observed to be correlated with the defined virulence categories or other meaningful data attributes (for varying k-values where appropriate or with DBSCAN).

Multilocus sequence types (STs) were assigned using the PubMLST database [49] with mlst v.2.19 on the assembled genomes and srst2 v0.2.0 using sequencing reads if not identified with MLST. The MLST composition of test and training data were confirmed to be of similar composition with no significant differences indicating appropriateness of training data for the classification task at hand. The *Klebsiella* genome cohort were additionally assessed for their virulence score and resistance profile using Kleborate v2.1.0 [31].

#### 4.8.3 Virulence class definitions

For this use case, we define four main virulence categories: high virulent, low virulent, colonization, and not virulent with the full heuristics described in Supplementary Table S2. The isolation source and host from a genome’s metadata were used to define its virulence category. Isolation sources that were more sterile were used to indicate an increase in virulence severity. For example, samples collected from more sterile sites e.g. blood, aspirate, cerebral spinal fluid, were considered high virulent. Samples collected from non-sterile body sites e.g., drainage, sputum, respiratory were considered low virulent. Colonization samples were defined as samples originating from rectal swabs, skin, or feces. Samples labeled as not virulent samples were those isolated from environmental or animal samples.

Two main classifications were modeled: 1. high versus low virulent and 2. virulent (including high and low) versus not virulent (including not virulent and colonization). In these comparisons, the more virulent category is considered Class1 (high and virulent, respectively) and the less virulent category is Class0 (low and not virulent, respectively).

#### 4.8.4 Feature description

For the *Klebsiella* genomes passing these metadata requirements, the respective domain architectures (DAs) were retrieved from the Functional Genomics Platform for each genome. These DAs are defined as pivot, co-pivot, or discovered DAs as previously described and are used as the feature set for a given genome with the number of DA occurrences per genome as its count value to define the ‘discovery inclusion’ feature set. This yielded 3,905 DA features across the genome cohort.

In order to assess accuracy of our discovery inclusion data, we also completed a comparison against a baseline *Klebsiella* feature set where only pivot proteins i.e. virulence seed data from VFDB were included. Neighbor and discovered domain architectures were not included. This resulted in 1,193 DA features for the baseline models.

#### 4.8.5 Feature selection

Feature selection was completed on the domain architecture count vectors for discovery inclusion feature set using two methods: 1. removal of features with zero variance and 2. a deduplication where one feature vector is selected to represent a group of features if there was greater than 99% correlation within the group. The number of features removed varied as a function of the number of features analyzed based on host preference and the classification model itself. Depending on the model and host type, 345 to 688 (on average, approximately 530) features were dropped from the total 3,905 features (Supplementary File S1).

We leveraged XGBoost [50] with no further feature scaling or feature extraction for both discovery inclusion and baseline models. SVM was explored with the discovery inclusion feature set and standard scaling from Python sklearn [51] and PCA were used for these models. Variance captured by PCA was tuned as a hyperparameter.

#### 4.8.6 Model generation

To train our machine learning models, we evaluated multiple permutations for accuracy and biological relevance:

- **Data Segmentation:** For all models, 90% of the genome cohort was used in training and 10% was used for testing.
- **Klebsiella Sub-selection**: *Klebsiella pneumoniae* (Kp) is the major cause of clinical *Klebsiella* infections and is, therefore, of greater interest from a public health perspective; however, the usage of non-Kp genomes can potentially allow for better generalizablity. To this end, we evaluated a sub-selection of the *Klebsiella* cohort and feature set for just *Klebsiella pneumoniae* genomes or for all *Klebsiella* genomes. This was varied in testing and training data.

- Two test genome sets were created using a fixed random seed with either Kp-only or all *Klebsiella* genomes.
- Likewise for training, two similar sets were created i.e., one with Kp-only or one with all *Klebsiella* genomes.
- **Model Selection**: Binary and multiclass models were generated using the virulence category definitions. However, multiclass models where each virulence category was treated as an independent class did not surpass chance performance due to dimensionality limitations and were not included in this work. Binary models of high versus low virulent and virulent versus not virulent as previously described in Methods Section 4.8.3 were therefore employed for the characterization of clinical *Klebsiella*. Positive unlabeled learning was also evaluated with the consideration that some *Klebsiella* infections initially may have been isolated from a low virulent isolation source but could progress as the infection increased in severity; however, positive unlabeled learning did not achieve significant accuracy and was not pursued further.
- **Host Selection:** Training sets were generated for only human hosts or all hosts.
- **Model Architecture Selection:** We relied on XGBoost and SVM as our model algorithms. Neural networks were evaluated but did not yield strong performance which is congruent with prior works as they are expected to underperform XGBoost and SVM when data size and diversity are limited e.g., less than 50K data points [52].

#### 4.8.7 Model tuning

Regularization of the models was performed, and regularization weight was tuned as a hyperparameter using sklearn [51] during model generation. Hyperparameter tuning was performed for all models using Bayes Search Cross-Validation in the Python scikit-optimize package with 5-fold crossvalidation and up to 100 iterations of Bayes Search. Models were trained with and without oversampled data.

#### 4.8.8 Model testing and validation

Model performance was evaluated by calculating F1-Score, precision, recall, and AUC for ROC using Python sklearn [51]. Additionally, models were evaluated for overfitting by generating learning curves as a function of the number of samples a model had processed.

In this use case, we worked with publicly available genomic data which comes with bias due to the increased need for sequencing of clinical and pathogen isolates; therefore, the data will naturally exhibit a class imbalance where the majority class (i.e. virulent) will have more observations. To ensure that the evaluation metric best represented the model performance without overemphasizing a single class, we used macro-averaged F1-Scores which treat both classes with equal weight opposed to F1-Score which would be biased toward the majority class. Precision and recall are reported for each class.

To assess the model performance across *Klebsiella* species, error analysis by MLST was performed. MLST were assigned as previously described (Section Methods 4.8.2). Of the 176 MLSTs analyzed, many were represented by one or only a few genome(s) in the test dataset and predominantly belonged to only one class. Therefore for error analysis, the *zero division* parameter in scikitlearn’s *classification matrix* function was set to *np.nan*. This allowed for differentiation between a true zero in the precision and recall calculation by ST versus an NA value representing unable to be detected when there were no samples for the class. The default behavior was used when performing model evaluation on the entire test set.

## Supplementary Information

### Supplementary Material

- Supplementary Figures S1-S8 and Supplementary Tables S1-S4
- Supplementary-File-S1-number-features-dropped-discovery-inclusion.csv
- Supplementary-File-S2-stacked-unique-features-per-class.txt

## Declarations

The findings and conclusions in this report are those of the authors and do not necessarily represent the views of the Centers for Disease Control and Prevention.

## Code Availability

Code relating to the genera agnostic virulence prediction method is available on GitHub https://github.com/IBM/omxware-getting-started/tree/master/Virulence-Analysis/methodology as well as the *Klebsiella* model generation and machine learning analysis https://github.com/IBM/omxware-getting-started/tree/master/Virulence-Analysis/machine_learning_use_case-Klebsiella.

## Availability of Data and Materials

Predicted virulence discoveries for all genera have been deposited to PrecisionFDA as a Spark warehouse https://precision.fda.gov/home/assets/file-Gj2qKX00ZqJppv3JjxPFxJkG-1. From the *Klebsiella* virulence prediction use case, the feature tables, models, relevant metadata, and classification outputs have also been open-sourced to PrecisionFDA https://precision.fda.gov/home/assets/file-Gj87zv00ZqJQ36gqppFz9QQF-1. Please see GitHub for complete documentation of data tables https://github.com/IBM/omxware-getting-started/tree/master/Virulence-Analysis/methodology.

## Notes

https://github.com/IBM/omxware-getting-started/tree/master/Virulence-Analysis/

https://precision.fda.gov/home/assets/file-Gj2qKX00ZqJppv3JjxPFxJkG-1

https://precision.fda.gov/home/assets/file-Gj87zv00ZqJQ36gqppFz9QQF-1

### Competing Interest Statement

The authors have declared no competing interest.

