## Supplementary Figures S1-S8 and Supplementary Tables S1-S4 for "Commensal or pathogen: computationally vectorizing microbial genomes for *de novo* risk assessment and virulence feature discovery in *Klebsiella pneumoniae*"

**Table S1:** FGP IPR Domain Summary

| IPR Type | # Distinct IPR | # Distinct Protein |
|---|---|---|
| ACTIVE_SITE | 129 | 969,187 |
| BINDING_SITE | 73 | 861,228 |
| CONSERVED_SITE | 606 | 6,301,098 |
| DOMAIN | 6,827 | 29,013,089 |
| FAMILY | 14,591 | 24,771,707 |
| HOMOLOGOUS_SUPERFAMILY | 2,569 | 30,900,236 |
| PTM | 12 | 250,448 |
| REPEAT | 236 | 912,019 |

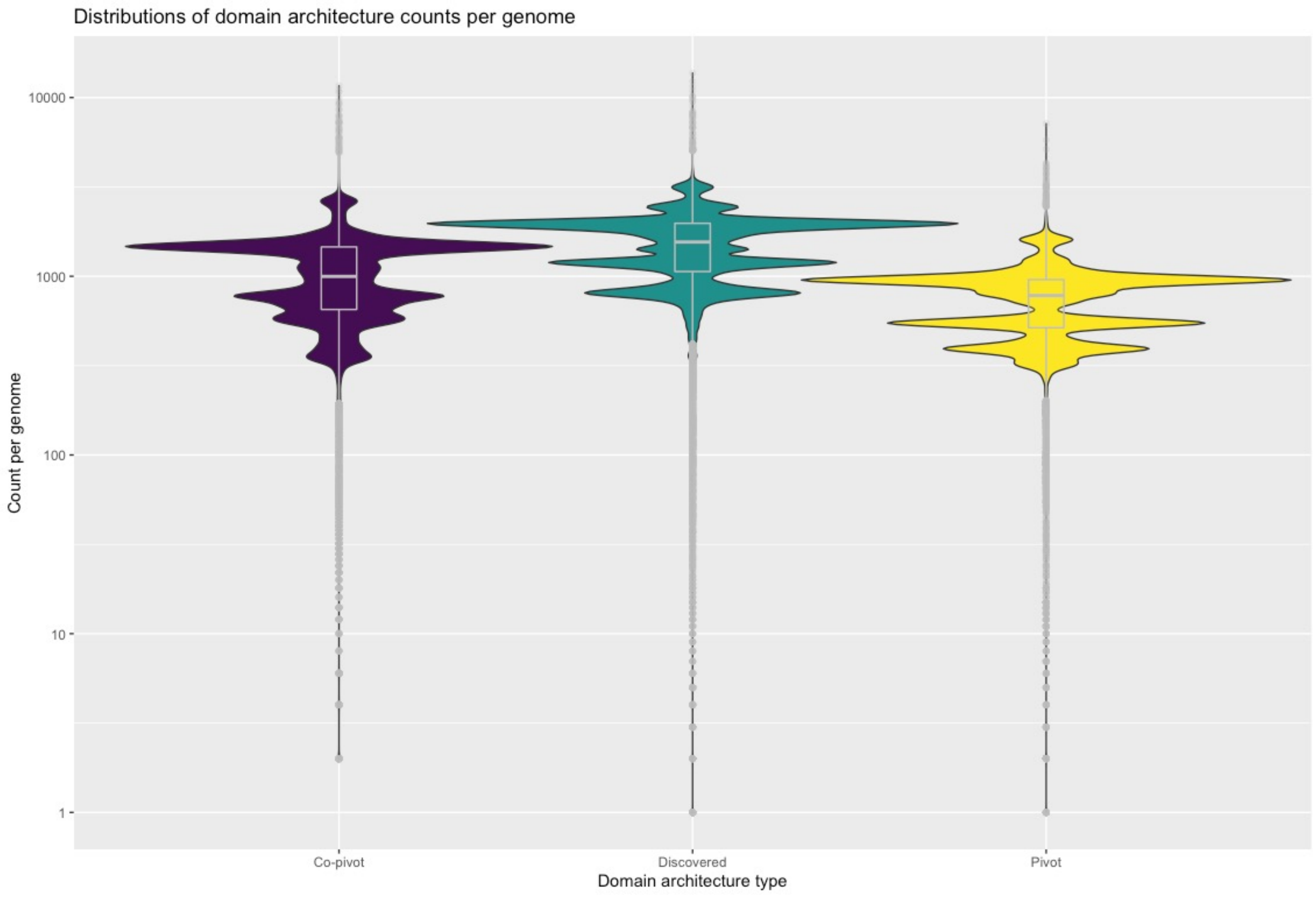


**Figure S1:** Distribution of domain architecture types per genome. The total count of unique co-pivot, discovered, or pivot domain architectures in a given genome is plotted for the entire genome corpus analyzed.

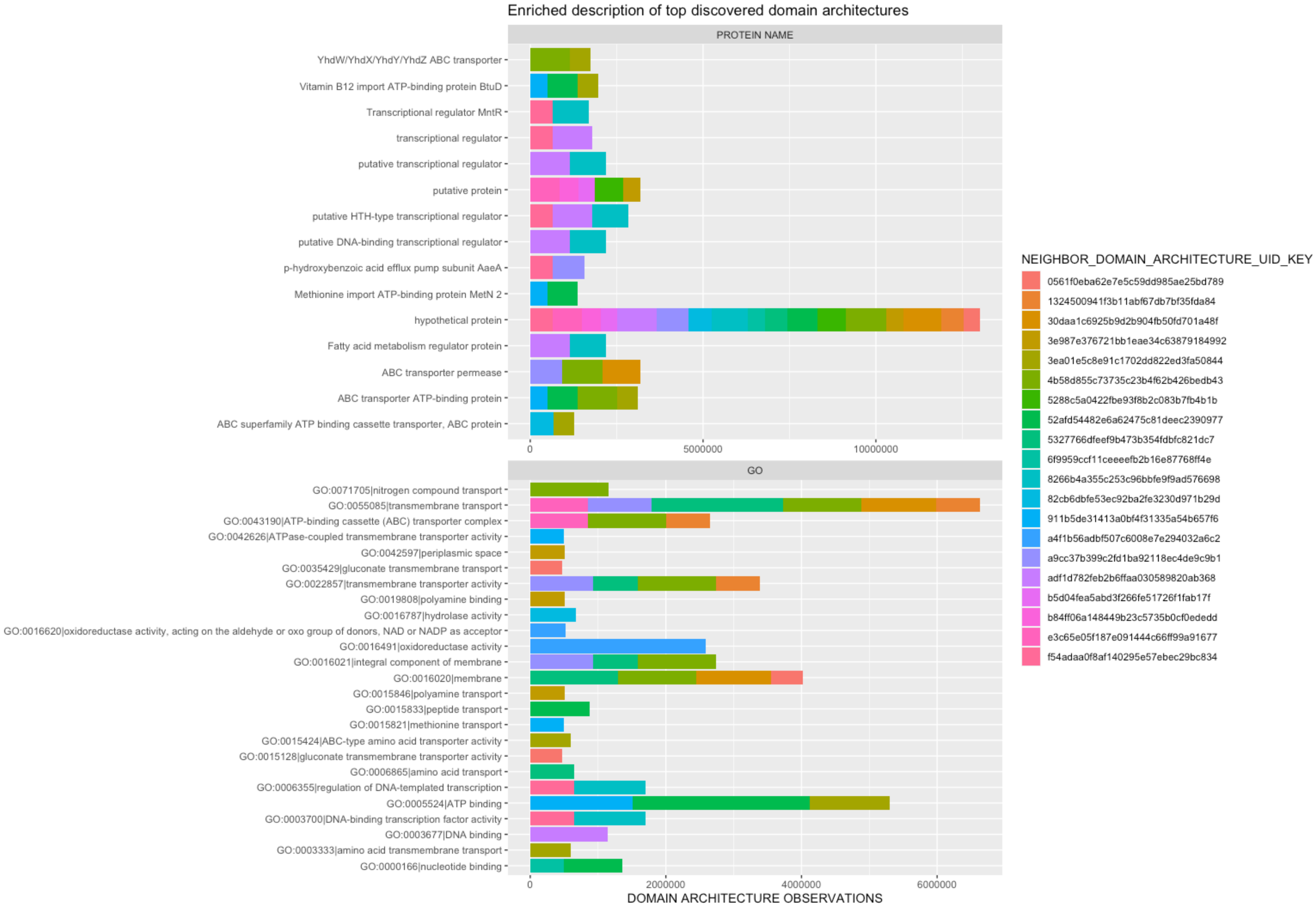


**Figure S2:** Top twenty discovered domains architectures (DA) by occurrence across all genera are described by the set of enriched protein names (protein names occurring for more than one top DA) in top panel and their respective GO terms (bottom panel).

A.

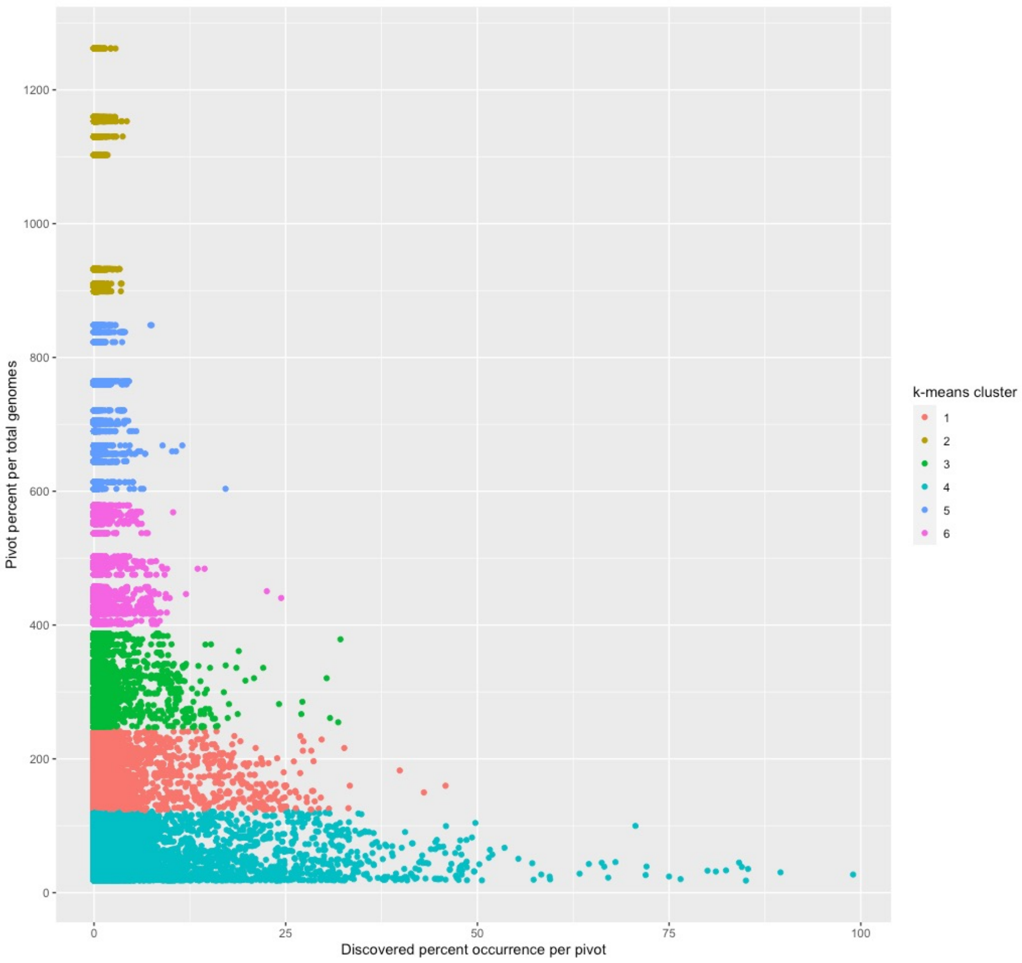


B.

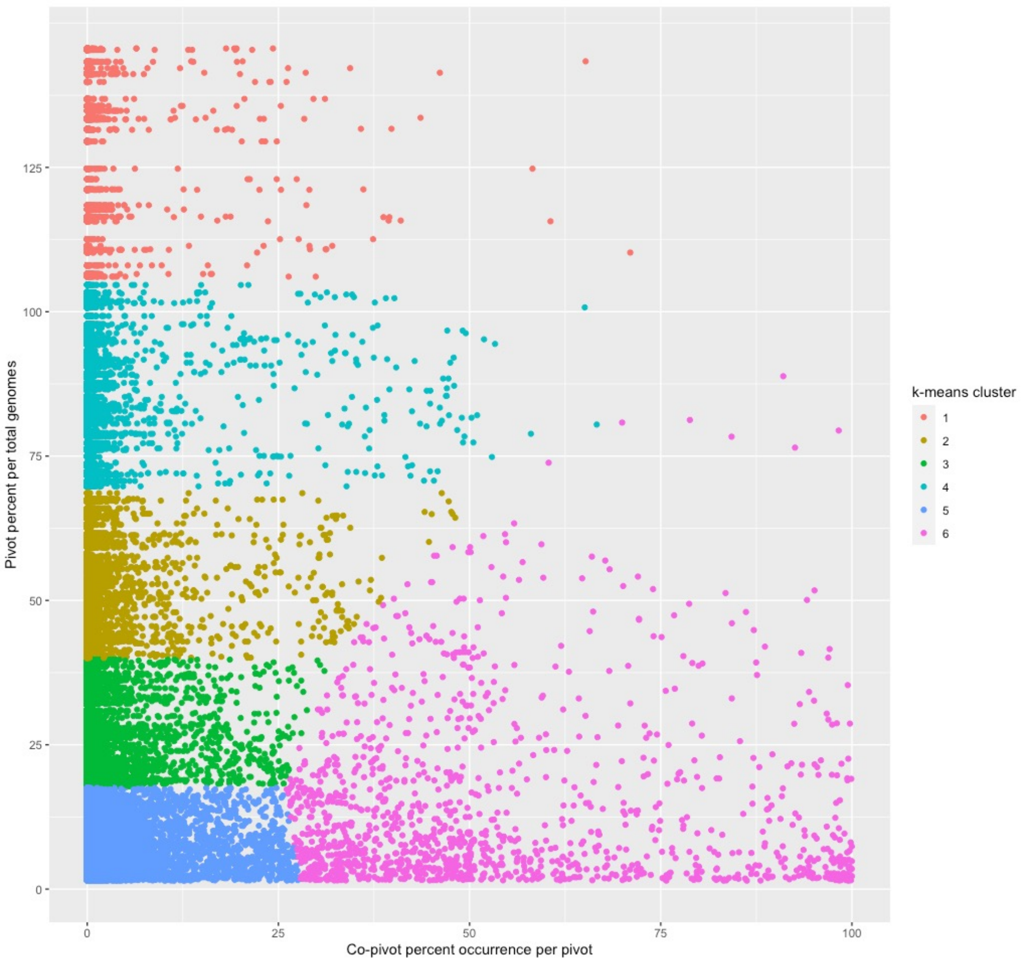


**Figure S3:** Occurrence of pivot:co-pivot and pivot:discovery domain architecture pairings across all genera are clustered using k-means. The percentage a pivot DA occurs across all genomes in a cohort is indicated in the y-axis as well as the percent that a discovered (A) or co-pivot (B) domain architecture occurs as its direct neighbor is shown.

### **Table S2:** Virulence Category Definitions

| Virulence Category | Definition |
|---|---|
| NOT_VIRULENT | Environmental and animal isolates--note that enviornmental isolates are not here differentiated by clinical or non-clinical, acknowledging clinical environmental isolates are likely human-associated and therefore may be at least as virulent as colonization |
| COLONIZATION | Rectal swab, skin, feces |
| LOW_VIRULENT | All clinical isolates from body sites or specimen sources that are from 'non-sterile body sites' (incl drainage, sputum, respiratory, wound, and urine) |
| HIGH_VIRULENT | Blood, aspirate, abscess, CSF, tissue |
| MISSING** | Missing, dash, empty, 'non-applicable', etc |
| AMBIGUOUS_CLINICAL** | Associated with healthcare but cannot differentiate virulence level--note that some body fluids appear here although they could be from even 'sterile body site', don't know how they were collected and whether indwelling drain may have become colonized |
| AMBIGUOUS_GENERAL** | Not well-associated with healthcare and cannot differentiate virulence level--note that isolates labled only by hospital name appear here |

***Denotes categories excluded from model creation due to insufficient information.*

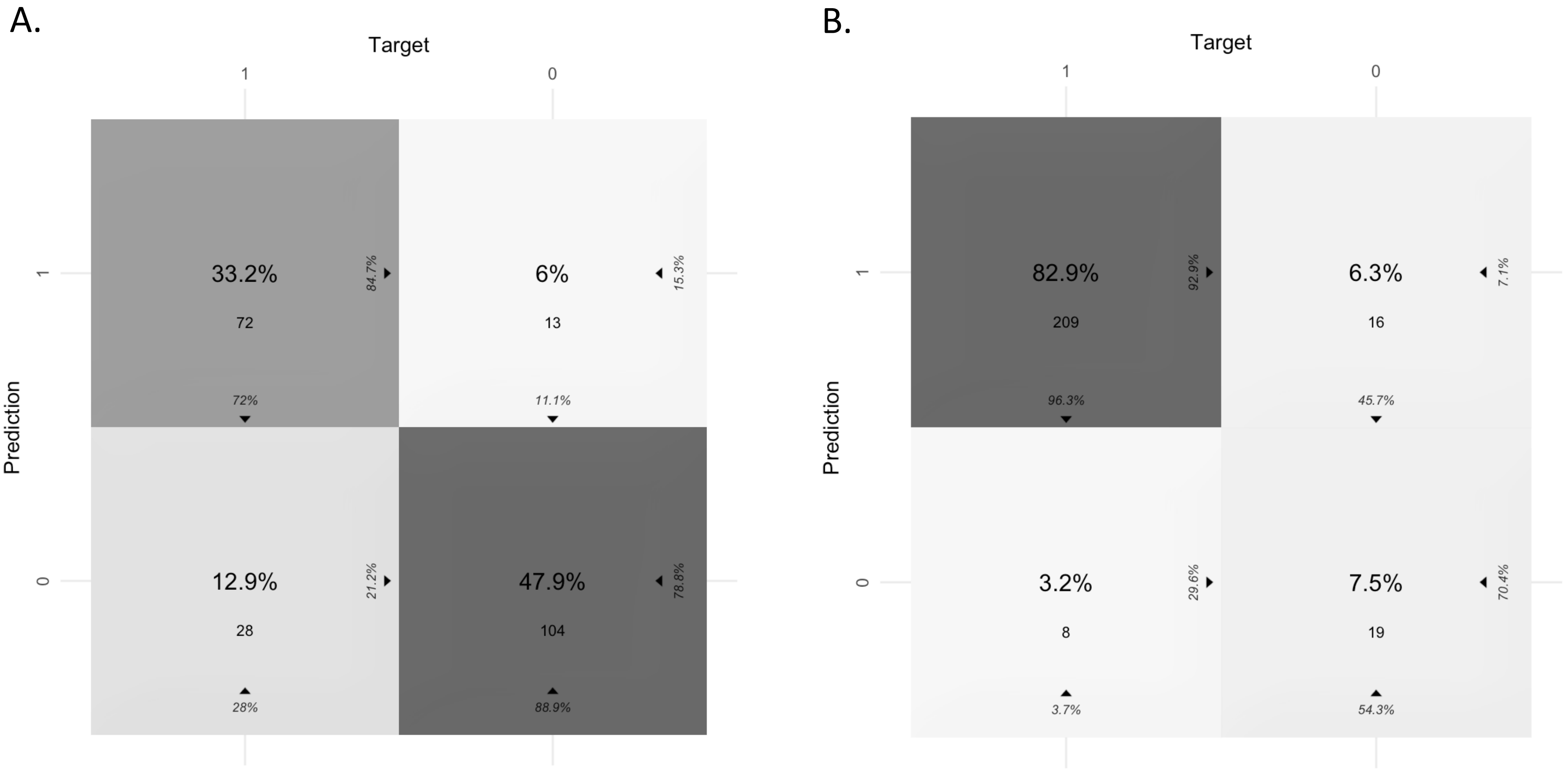


**Figure S4:** Confusion matrices are shown for the best performing models of each classification task: high versus low virulence (A) and virulent versus not virulent (B).

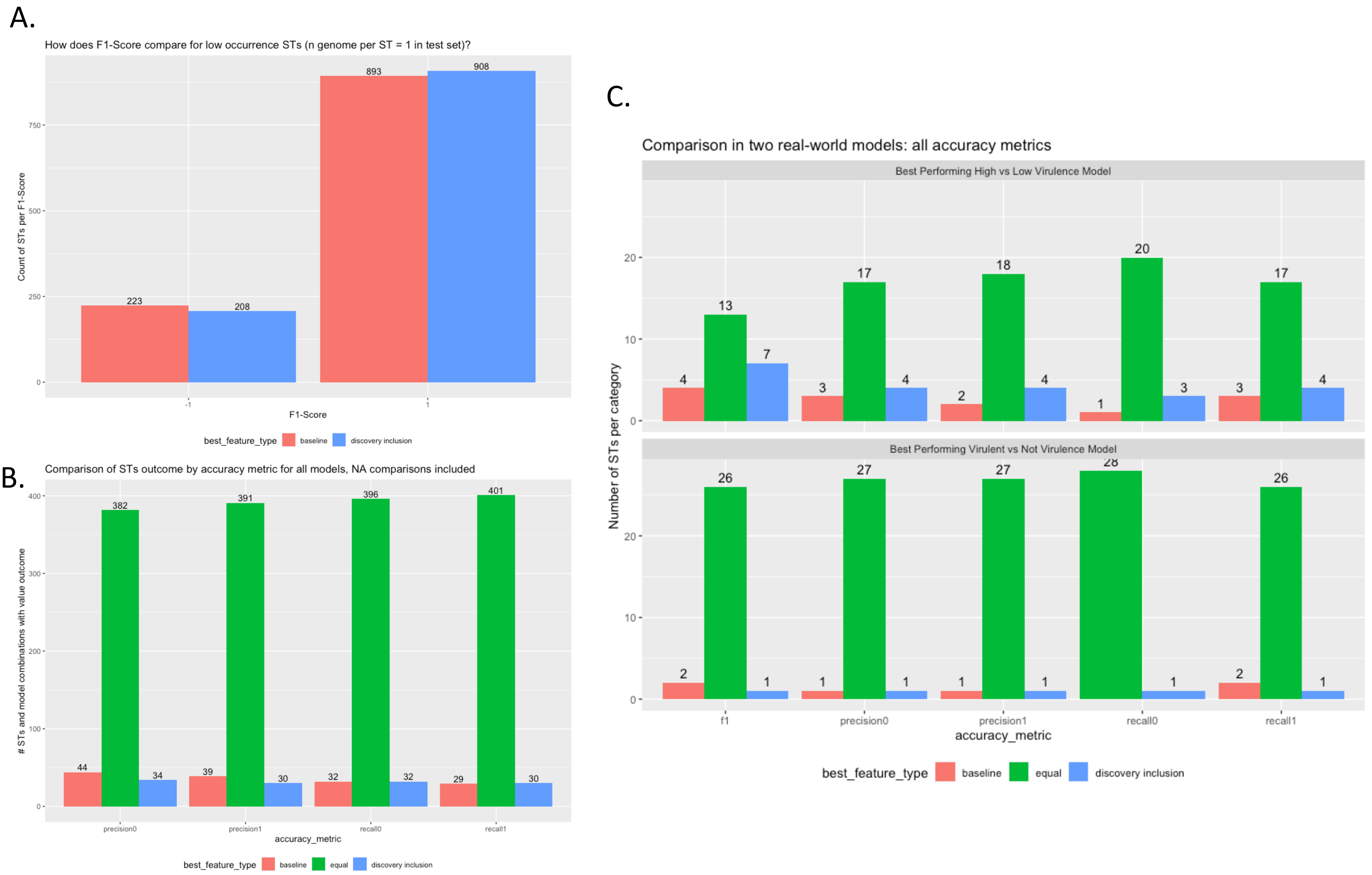


**Figure S5:** Error analysis by MLST are indicated for baseline and discovery inclusion feature sets. The F1-Score is compared for STs with one genome in the test set (A) where -1 is a representation of F1-Score = NaN. The precision and recall values for all other STs are evaluated for <u>all models</u> for which feature set is better performing or if the accuracy metrics indicate the feature sets yield equivalent results and the counts are shown (B). The best performing models of each classification task are shown in (C) with the feature type that yielded the highest accuracy.

**Table S3:** MLST Diversity by Virulence Category

| VIRULENCE CATEGORY | # distinct MLST | # samples | MLST diversity per samples |
|---|---|---|---|
| NOT_VIRULENT | 54 | 102 | **0.53** |
| COLONIZATION | 78 | 250 | **0.31** |
| HIGH_VIRULENT | 232 | 1089 | **0.21** |
| LOW_VIRULENT | 195 | 1361 | **0.14** |

**Table S4:** VFDB IPR Summary

| IPR Type | # Distinct IPR | # Distinct Protein |
|---|---|---|
| ACTIVE_SITE | 30 | 556 |
| BINDING_SITE | 19 | 247 |
| CONSERVED_SITE | 81 | 2,781 |
| DOMAIN | 923 | 13,919 |
| FAMILY | 1,101 | 13,851 |
| HOMOLOGOUS_SUPERFAMILY | 541 | 12,971 |
| PTM | 5 | 675 |
| REPEAT | 59 | 830 |

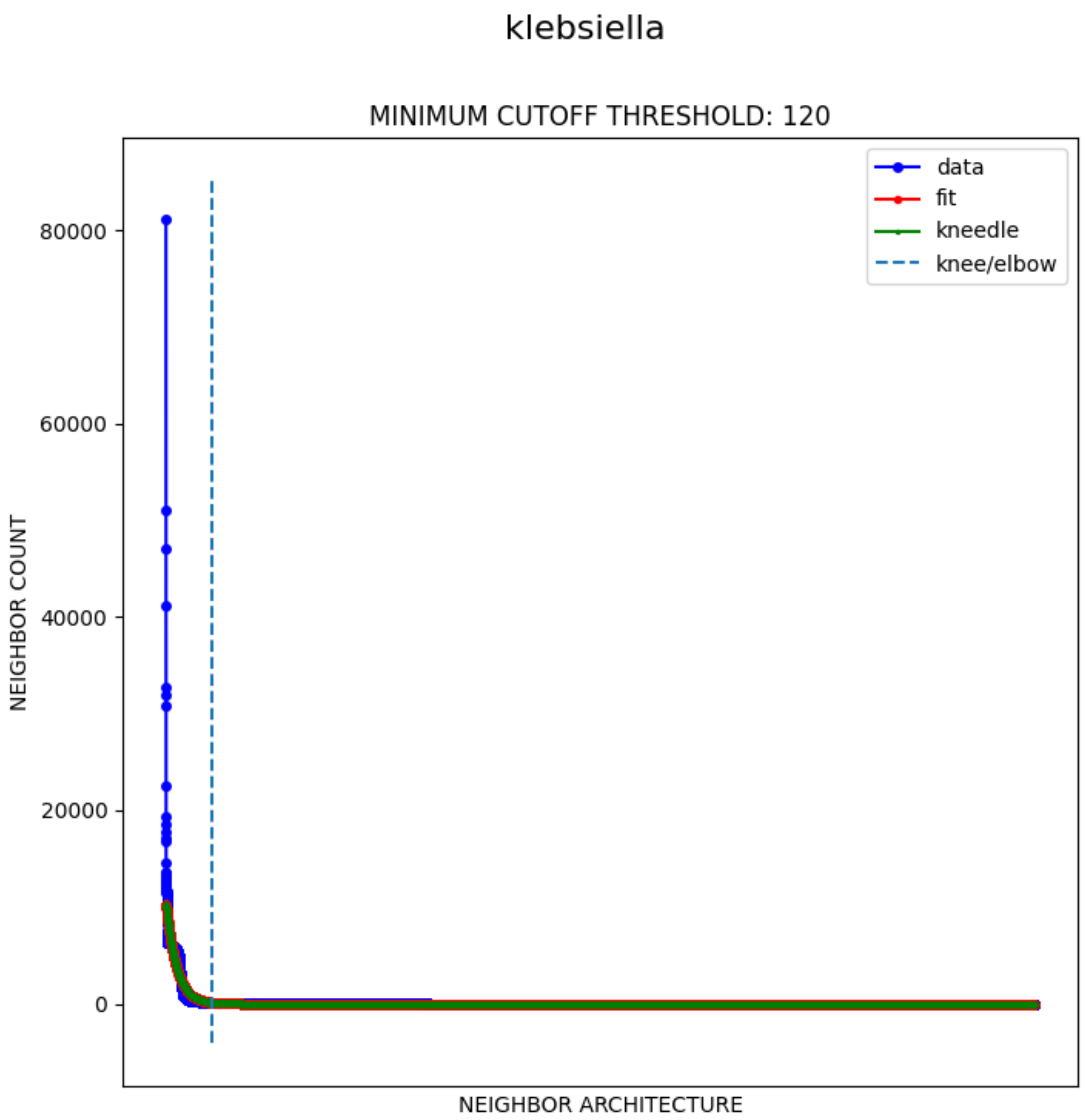


**Figure S6:** Example usage of computing minimum cutoff threshold for *Klebsiella* using Kneedle algorithm where neighbor count is observed count within that genera and neighbor architecture is the sorted domain architecture hash.

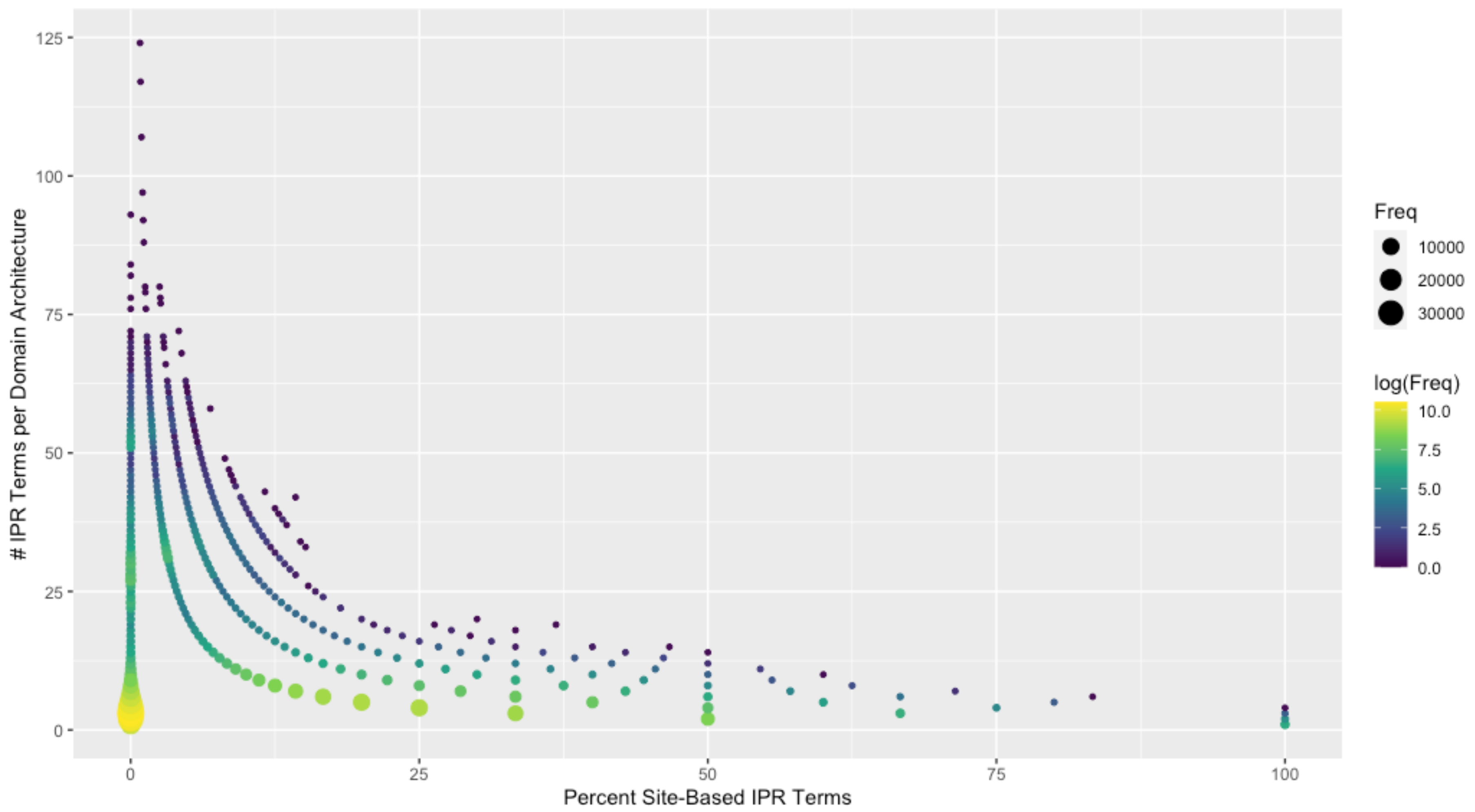


**Figure S7:** Assessment of Site-based IPR Terms in Domain Architectures. All domain architectures from seed data collection were assessed for the occurrence of site-based terms (ACTIVE SITE, BINDING SITE, CONSERVED SITE, PTM, or REPEAT) where the total number of IPR terms is contrasted against the percentage of those terms that are site-based. The frequency for that numerical combination is provided in color and circle size.

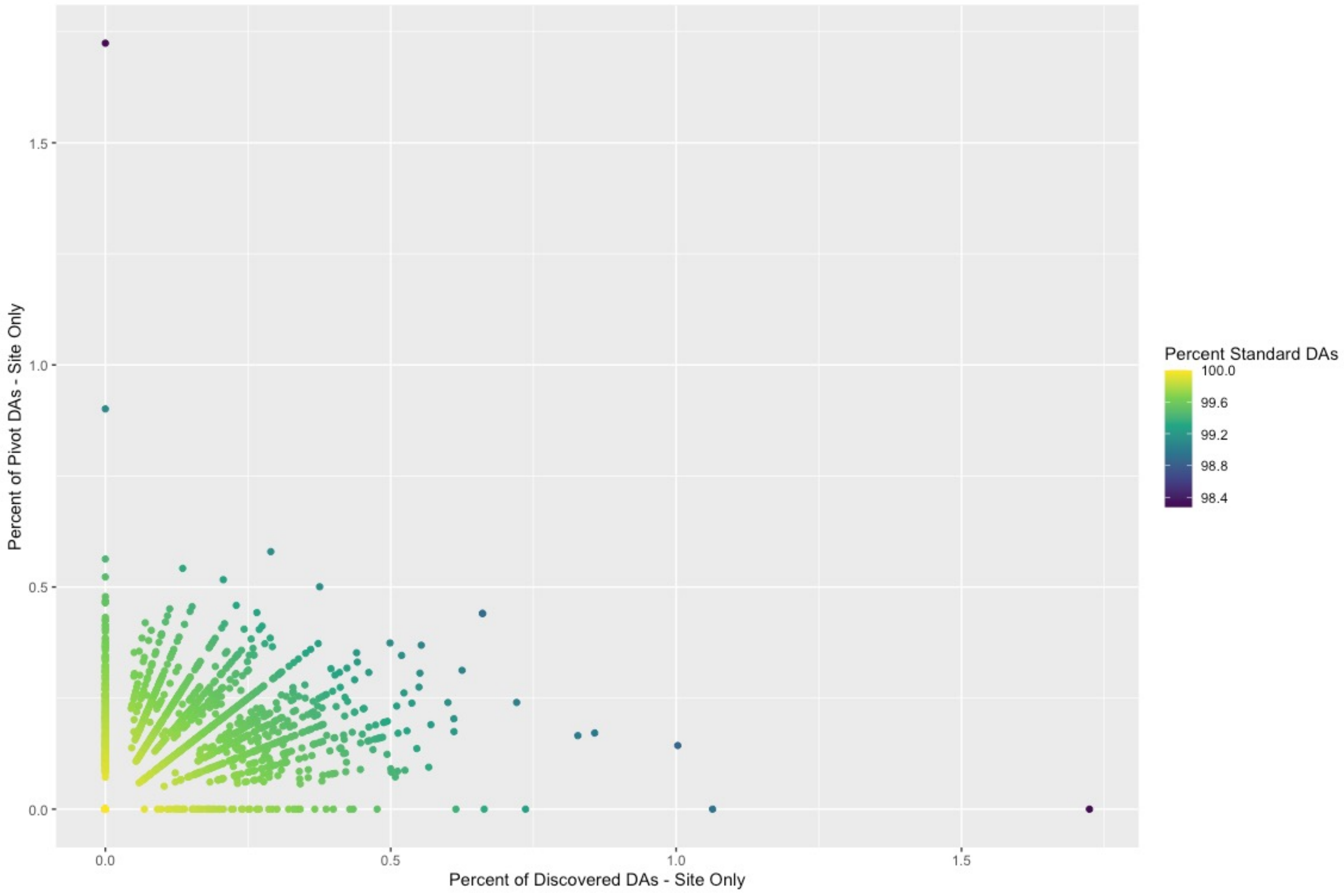


**Figure S8:** Assessment of Site-based IPR Terms in Discovered Domain Architectures. For each genera, the percentage of domain architectures comprised of all site-based terms is contrasted for discovered DAs and pivot DAs. The percentage of non-site-based terms i.e. standard intermixed domains architectures is indicated in color. Domain architectures included in this plot are those above the thresholds applied in kneedle analysis.
